# TIMPANI: Target-Informed Pretraining for Retention-Time Prediction across Chromatographic Conditions

**DOI:** 10.1101/2025.06.26.661631

**Authors:** Yuhui Hong, Haixu Tang

**Author notes:** Yuhui Hong and Haixu Tang. 2027. TIMPANI: Target-Informed Pretraining for Retention-Time Prediction across Chromatographic Conditions. In *Proceedings of the 33rd ACM SIGKDD Conference on Knowledge Discovery and Data Mining V*.*1 (KDD ‘27), August 01–05, 2027, San Jose, CA, USA*. ACM, New York, NY, USA, 12 pages. https://doi.org/10.1145/3862868.3871680.

## Abstract

Retention time (RT) in liquid chromatography is a key signal for small-molecule identification, yet effective modeling remains challenging because each chromatographic condition contains only a limited number of labeled compounds, and retention orders sub-stantially differ across conditions, resulting in poor cross-condition transfer. To investigate transferability, we first show that transfer performance is governed primarily by the distance between chromatographic conditions rather than by molecular overlap (Pearson *r* = −0.65 vs. −0.14). Because well-labeled conditions are rarely close to a given target, single-source transfer is unreliable, motivating a pre-training strategy that aggregates data across diverse conditions while adaptively emphasizing those most similar to the target. Guided by this insight, we present TIMPANI (*Target-Informed Pretraining*). Unlike conventional transfer learning or meta-learning, where the target remains unseen until adaptation, TIMPANI assumes that a small set of measurements from the target condition is available during pre-training. It meta-learns an initialization across upstream conditions, using these measurements to anchor pre-training and steer the pool toward the conditions most relevant to the target, before fine-tuning with the encoder frozen. On data-scarce conditions where conventional transfer learning performs poorly, TIMPANI significantly improves prediction accuracy over it (mean *R*^2^ from 0.60 to 0.68, *p* = 0.02), while three recent RT baselines fail to converge in this low-data regime. Across all 36 eligible RepoRT datasets, it is no worse than conventional transfer learning on average (*R*^2^ 0.84 vs. 0.83), with the gains concentrated on such difficult conditions. Without per-dataset chromatographic metadata (only the RP/HILIC mode label selects the source pool), it performs comparably to the only stable external baseline, a recent metadata-conditioned graph transformer. Our controls attribute this result to target fine-tuning, which compensates for the missing metadata. TIMPANI thus makes accurate, condition-specific RT prediction practical for laboratories with only a small in-house dataset, exploiting abundant public RT data.

## 1 Introduction

Liquid chromatography (LC) is a cornerstone technique for separating and identifying compounds in complex mixtures [7], with applications across metabolomics [6], environmental science [5], and pharmaceutical research [23]. Retention time, the time a compound takes to elute from the LC column, is a key signal for small-molecule identification and has been modeled for decades [13] with progressively richer molecular representations, from classical descriptors [3] to molecular graphs [31] and three-dimensional conformers [14, 15]. Deep learning performs especially well on large benchmarks collected under a single standardized condition, such as METLIN-SMRT [8].

In practice, however, LC conditions vary widely in stationary phase, mobile phase, flow rate, and temperature to resolve different compound classes. The RepoRT collection [19] shows the effect, with retention times correlating within a chromatographic mode but anti-correlating across modes. This diversity is hard for deep learning, which needs large, condition-specific training sets, yet most conditions yield few labeled compounds because data acquisition is slow and costly. Models therefore struggle to generalize across conditions, especially with few labeled compounds [16, 20].

A common remedy is transfer learning (TL), which pre-trains on a single data-rich upstream task (e.g., a standard LC condition), then fine-tunes on the data-scarce downstream task [20]. But a single broad upstream source often gives a poor initialization for a specific condition, because its representations miss system-specific detail. A model tuned to reversed-phase chromatography, for instance, transfers poorly to normal phase, and even within reversed phase, column chemistry, additives, or temperature shift retention substantially. Each chromatographic system thus calls for a more targeted transfer strategy.

Inspired by gradient-based meta-learning [11, 25], we propose **TIMPANI** (**T**arget-**I**nfor**m**ed **P**retr**a**i**ni**ng). TIMPANI pre-trains a shared initialization over a pool of upstream tasks (distinct LC conditions) while anchoring on the target (the new condition, for which few samples are available), then adapts by fine-tuning the prediction head with the encoder frozen. Unlike TL, the target’s own training data enter during pre-training, yielding an initialization from which the target optimum is reached in a few samples, not merely one that is fine-tunable in principle. Unlike standard meta-learning, which seeks one initialization that adapts to many unseen tasks, TIMPANI pre-trains for one known target condition at a time, in a transductive setting. Because no single upstream condition fits every target, pre-training draws on the whole pool, with the target’s gradient shaping the initialization at every step. TIMPANI is backbone-agnostic, and when chromatographic meta-data are recorded, an optional module conditions the predictor on the experimental setup. Like a timpani tuned to its target pitch, TIMPANI tunes pre-training to its target task (Figure 1).

**Figure 1:**
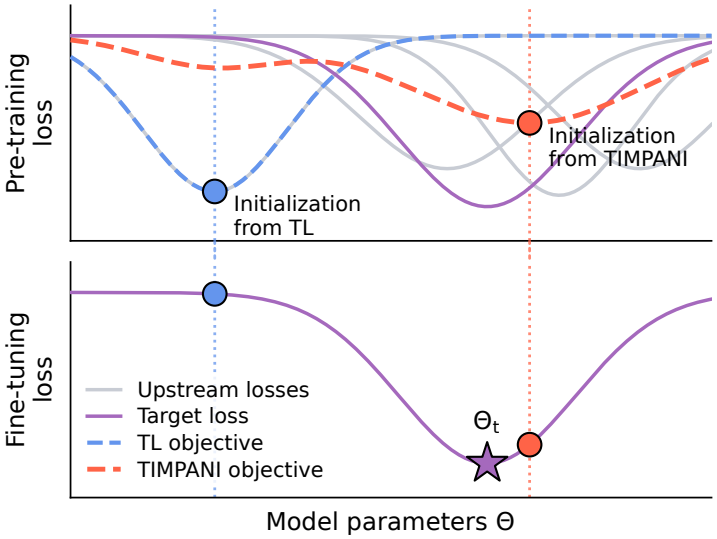
Target-informed pre-training initializes the model near the target optimum, whereas conventional transfer learning does not. Solid curves denote the loss landscape, and dashed curves indicate the objective optimized by each method. The top and bottom panels show the pre-training and fine-tuning losses, respectively, along a one-dimensional slice of the shared parameter space, Θ. Here, Θ_*t*_ denotes the optimal parameters for the target task.

In this paper, we address two research questions. First, does anchoring pre-training on the target improve on conventional TL, particularly where TL’s accuracy is poor? Second, once the model is fine-tuned on the target, how much do the recorded chromatographic metadata contribute to performance? We evaluate TIMPANI on the RepoRT datasets where conventional TL fails to reach satisfactory accuracy (*R*^2^ < 0.8), under 5-fold cross-validation on a Graph Isomorphism Network (GIN) backbone, and compare against four recently developed RT prediction methods, including the metadata-aware Graphormer-RT [18, 22, 27, 29].

Our contributions are summarized as follows:

- **A diagnosis of when TL fails**. Transfer performance tracks chromatographic-condition distance to the source (*r* = −0.65) more than molecular overlap, so pre-training must pool diverse conditions.
- **A target-informed pre-training strategy**. TIMPANI meta-learns a target-specific initialization over a pool of source conditions, using the target to anchor pre-training and re-weight sources by gradient alignment, then fine-tunes on the target, letting a lab build a condition-specific model from a small in-house dataset plus public data.
- **Gains without per-dataset chromatographic metadata**. Using only the RP/HILIC mode label, not per-dataset meta-data (column, mobile phase, gradient), TIMPANI raises mean *R*^2^ over TL from 0.60 to 0.68 (*p* = 0.02) and lowers MedAE from 0.92 to 0.77 min on ten *TL-difficult* RepoRT datasets, and is no worse than TL over all 36 eligible datasets (0.84 vs. 0.83). The gain holds on fresh fold partitions and scaffold-disjoint splits (Section 5.2). TIMPANI performs comparably to Graphormer-RT, the only stable one of four recent base-lines in this low-data regime, and an ablation attributes the gain to gradient-alignment re-weighting (*p* = 0.002).
- **A finding that target fine-tuning compensates for missing metadata**. Removing the target’s recorded metadata barely affects Graphormer-RT (*p* = 0.56), and resampling it does not hurt either: target fine-tuning compensates for the missing fields, which explains why a metadata-free model can perform comparably to a metadata-aware one.

## 2 Condition Distance Contributes More Than Molecular Overlap to Transfer

LC retention data are scarce and condition-diverse: most RepoRT datasets carry only 100–300 labeled compounds (Appendix A.1), and their conditions span reversed-phase (RP) and hydrophilic-interaction (HILIC) modes whose retention orders differ sharply (Figure 2d). On such data, is TL enough? Following Kwon et al. [20], who found that a GIN pre-trained on METLIN-SMRT and fine-tuned per target outperforms other graph neural network (GNN) architectures on most RepoRT datasets, we reproduce this setup under 5-fold cross-validation across the 38 datasets with enough eluting compounds to train on (>100; curation in Section 4). This baseline pre-trains on the single reversed-phase SMRT source and fine-tunes on each target, irrespective of its chromatographic mode.

**Figure 2:**
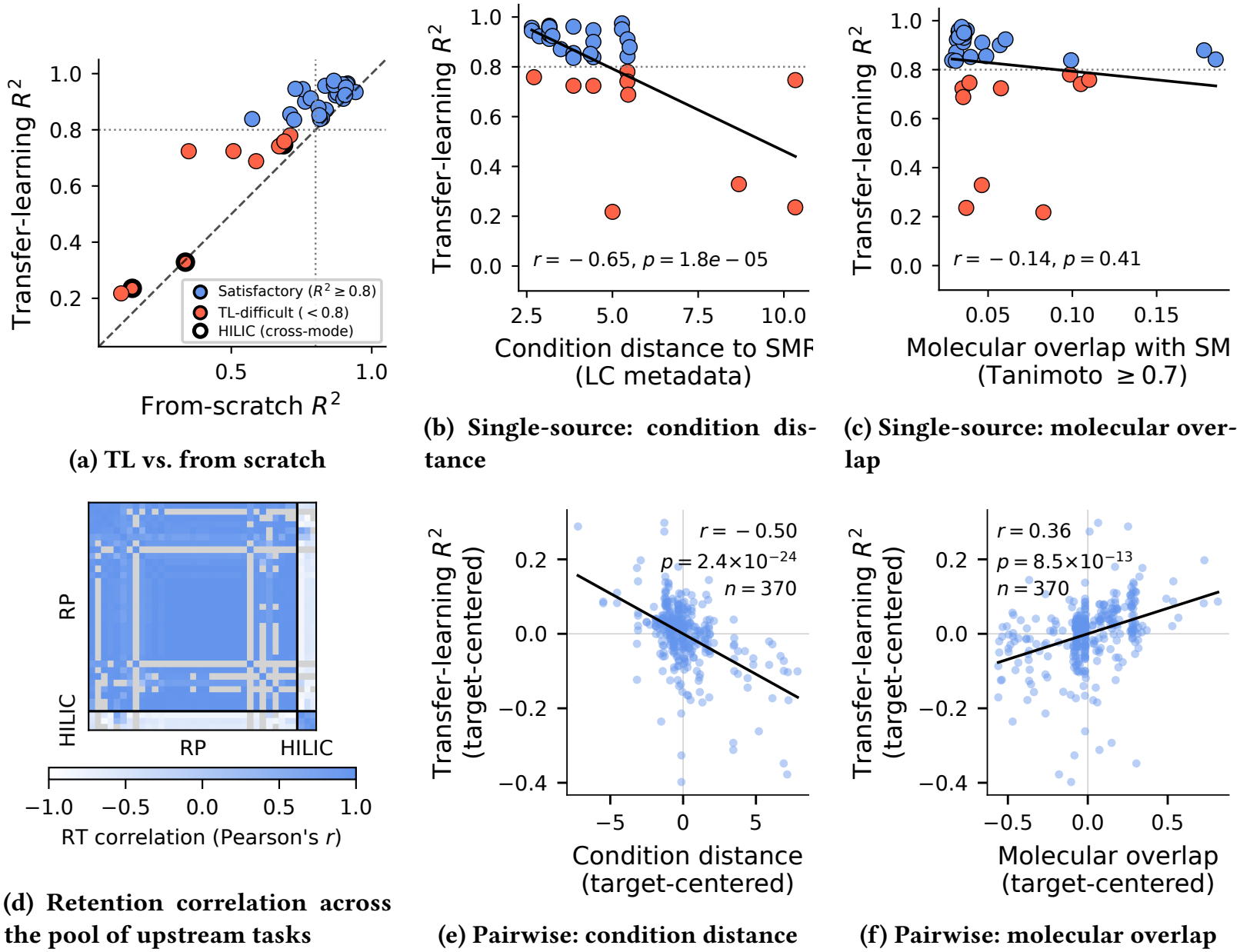
Single-source TL performance against two source-to-target properties, chromatographic-condition distance and molecular overlap, from one fixed source (top) and across all source-to-target pairs (bottom); see Section 2 for details. Top (single METLIN-SMRT source): (a) TL vs. from-scratch *R*^2^ per screened dataset: TL-difficult targets (TL *R*^2^ < 0.8) in red, HILIC (cross-mode) targets outlined; (b, c) TL *R*^2^ against condition distance to SMRT and against molecular overlap with SMRT (Tanimoto ≥ 0.7). Bottom: (d) pairwise retention-time correlation across the pool of upstream tasks, ordered by mode (RP, HILIC; gray: fewer than five shared compounds); (e, f) transfer *R*^2^, target-centered, against source-to-target condition distance and molecular overlap across 370 single-source-TL pairs (37 sources per target, excluding self-transfer). Scatter panels annotate Pearson’s *r*.

**Figure 3:**
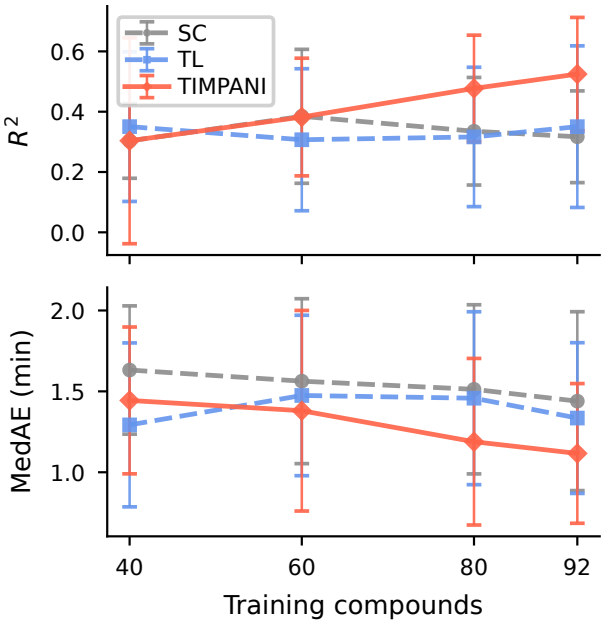
Data efficiency on RepoRT 0183 (GIN, 5-fold CV): *R*^2^ (top) and MedAE (bottom) for SC, TL, and TIMPANI as the target training set grows from 40 to about 92 compounds (the full fold). TIMPANI pulls ahead once the target reaches roughly 60 compounds, while TL and SC plateau; below that the three are within noise. As in the benchmark, TL uses METLIN-SMRT and TIMPANI the mode-matched pool.

Across this benchmark, TL consistently improves RT prediction over training from scratch (SC), most clearly where SC is weakest (Figure 2a, points above the diagonal). Yet ten datasets remain below *R*^2^ = 0.8 even with TL. We call these *TL-difficult* and evaluate TIMPANI on them (Figure 2a, red points). The *R*^2^ = 0.8 cutoff is pragmatic, as TL performance is continuous rather than bimodal, so our conclusions do not depend on its exact value. TIMPANI’s improvement over TL grows as TL performance falls and is negligible where TL already succeeds (Table 1, Figure 5).

**Table 1:** TL vs. TIMPANI (GIN, 5-fold CV, seed 42) on all 36 eligible RepoRT datasets and subsets. TL-difficult: TL *R*^2^ < 0.8. Far/near and small/large: extreme thirds by condition distance to METLIN-SMRT and by size. Wins: datasets where TIMPANI has the higher *R*^2^. *p*: paired Wilcoxon on per-dataset means, unadjusted (–: not tested, *n* < 6).

|  |  | Mean *R*<sup>2</sup> |  | TIMPANI vs. TL |  |  |
|---|---|---|---|---|---|---|
| Datasets | *n* | TL | TIMPANI | Δ*R*<sup>2</sup> | Wins | *p* |
| All eligible | 36 | 0.826 | 0.837 | +0.011 | 16/36 | 0.98 |
| TL-difficult | 10 | 0.597 | 0.676 | +0.080 | 8/10 | 0.020 |
| Other | 26 | 0.914 | 0.899 | −0.015 | 8/26 | 0.031 |
| Far | 12 | 0.693 | 0.746 | +0.052 | 7/12 | 0.18 |
| Near | 12 | 0.929 | 0.926 | −0.003 | 5/12 | 0.57 |
| HILIC | 3 | 0.437 | 0.627 | +0.190 | 3/3 | – |
| RP | 33 | 0.861 | 0.856 | −0.005 | 13/33 | 0.36 |
| Small | 12 | 0.716 | 0.771 | +0.055 | 7/12 | 0.15 |
| Large | 12 | 0.824 | 0.801 | −0.023 | 4/12 | 0.27 |

**Figure 4:**
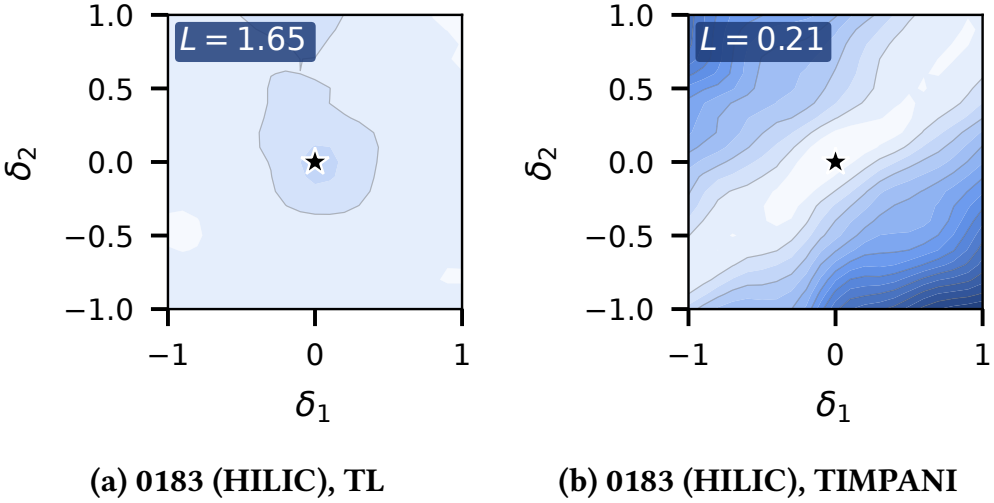
Downstream loss landscapes around the TL and TIMPANI initializations (GIN) on report 0183 (HILIC), following Li et al. [21]. The two panels share one color scale (lighter = lower loss; the black star marks the initialization). The TIMPANI basin is lower than TL’s, and the other targets show the same pattern (Section 6.3).

**Figure 5:**
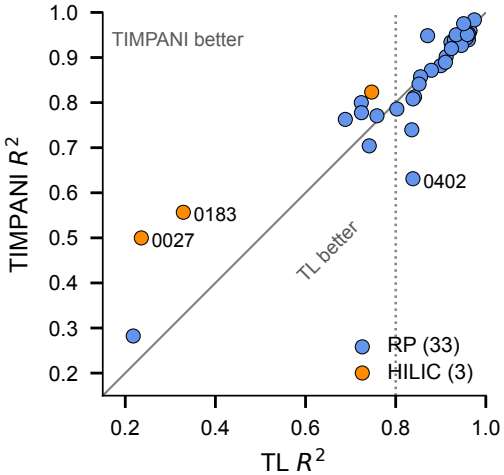
TL vs. TIMPANI *R*^2^ on each of the 36 eligible RepoRT datasets of Table 1, colored by mode. Points above the diagonal favor TIMPANI, and the dotted line marks TL *R*^2^ = 0.8. Per-dataset values are released with the code.

What makes a target TL-difficult? We relate TL from METLIN-SMRT (RepoRT dataset 0186) to two properties of each target: its chromatographic-condition distance to SMRT (from LC metadata such as column, gradient, eluent, and flow) and its molecular overlap with SMRT (the fraction of compounds with a close analog, Tani-moto ≥ 0.7 [2]). Both featurizations are defined in Appendix A.2. Transfer falls steeply with condition distance (Figure 2b; *r* = −0.65) but appears nearly flat against molecular overlap under this single source (Figure 2c; *r* = −0.14), a flatness we revisit below once the source is varied.

Condition distance tracks the mode structure of retention, which is strongly correlated within a chromatographic mode and anti-correlated across it (Figure 2d; mean *r* = 0.84/0.78 within RP/HILIC, −0.57 across). The mismatch is starkest across modes, where reversed-phase SMRT fails on all three HILIC targets (Figure 2a). But it is not only a mode effect. Seven of the ten TL-difficult datasets are themselves reversed-phase, so within-mode condition diversity defeats TL too (Appendix A.1).

A single fixed source, however, cannot reveal how a source’s chemistry affects transfer. We therefore varied it, running TL to each of the ten targets from every one of the 38 screened datasets except itself (37 sources per target; 370 pairs). After centering each target’s transfer *R*^2^ to remove target difficulty, both properties predict transfer, largely independently. Condition distance remains primary (Figure 2e; *r* = −0.50, *p* < 10^−23^), and molecular overlap emerges as a clear secondary driver (Figure 2f; *r* = +0.36, *p* < 10^−12^). It appeared flat under SMRT only because that single, chemically broad source has near-constant overlap across targets. With both in one model, each stays significant (*β* = −0.44 vs. +0.27). TIM-PANI exploits this independent predictiveness, re-weighting the pool along both axes.

No single source is chromatographically compatible with every target, and each contains only 100–300 labeled compounds, so effective pre-training must pool multiple conditions. For each target, we build a mode-matched pool of same-mode RepoRT datasets, excluding the target itself and any dataset generated under the same chromatographic condition (Section 4). Pooling is not itself novel. The metadata-aware Graphormer-RT [27], our strongest baseline, also pre-trains on many RepoRT datasets, but it relies solely on upstream data, and its mode-level grouping is coarse, since conditions vary substantially within a mode. TIMPANI instead uses the target’s training data to anchor pre-training and re-weight the pool toward the conditions most relevant to the target. Because this changes both the pre-training data (single source to a pool) and the optimization strategy, we evaluate them separately. The Graphormer-RT comparison (Section 5.3) holds the pool fixed, while the ablation (Section 6.1) isolates pooling, target anchoring, and gradient-based re-weighting.

## 3 The TIMPANI Framework

The key idea behind TIMPANI is that the target chromatographic condition is known at training time. We exploit it by meta-learning a shared initialization over a pool of upstream tasks, using the target’s limited training data to anchor pre-training and re-weight the pool toward the conditions most relevant to the target (Section 3.1). Meta-learning rates follow convergence theory (Section 3.2), so the pool can compensate for scarce target labels. The resulting model is then fine-tuned by updating only the prediction head, with the meta-learned encoder frozen. TIMPANI is backbone-agnostic and can optionally encode recorded chromatographic metadata through a conditioning module (Appendix A.3).

### 3.1 Target-Informed Pretraining

Let *f*_*θ*_ denote a molecular encoder, *h*_*ψ*_ a prediction head, and (optionally) *z* a per-task condition context, giving a predictor *F*(*x*; *θ, ψ*, *z*). We meta-learn an initialization of the shared parameters Θ = {*θ, ψ*} over the upstream pool {*J*_i_} using first-order, ANIL-style modelagnostic meta-learning [11, 25]. Treating each dataset as a task, we split its labeled compounds into a *support* set *S* and a disjoint *query* set *Q*; the *inner loop* adapts only the head (and the optional context) on the support, holding the encoder fixed:

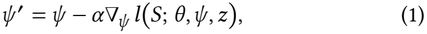

and the *outer loop* updates Θ on the query loss evaluated at the adapted head, with inner learning rate *α* and outer learning rate *β*.

The initialization is made target-informed by anchoring on the target. Crucially, the upstream pool {*J*_i_} deliberately excludes the target (and its same-condition replicates; Section 4); rather than letting the target appear occasionally as one sampled task among many, its training fold contributes a query gradient at every outer step with fixed weight, alongside a freshly-sampled batch of upstream tasks, and a single update is taken,

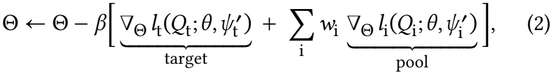

so the initialization is shaped by both the target and the pool at every step (Algorithm 1). Each source i is scaled by a weight *w*_i_ ∈ [0, 1] measuring how well its outer gradient aligns with the target’s. We define the alignment on two parameter groups. Partition the shared parameters Θ into the encoder Θ_enc_ (the molecular axis, which maps structure to a representation) and the condition context Θ_cond_ (the chromatographic-condition axis, the optional metadata module of Appendix A.3). Let *g*_i_ be source i’s outer (query) gradient and 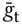 a running exponential-moving-average (EMA) reference of the target’s outer gradient, refreshed by the target anchor at every step. Writing *g*|_*G*_ for the sub-vector on group *G* and 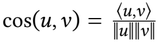, the per-axis cosines are

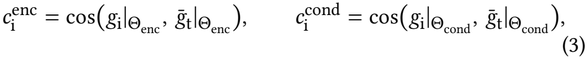

and, with the soft map *s*(*c*) = (1+*c*)/2 (anti-aligned → 0, orthogonal → 0.5, aligned → 1),

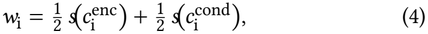

sources whose gradients point with the target contribute more to the pre-training. On a metadata-free backbone, there is no condition context (Θ_cond_ = *∅*); the condition axis is then undefined and set neutral, 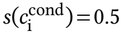; as a result, the weight is driven solely by the molecular axis and lies in [0.25, 0.75], whereas the full range [0, 1] applies only with the condition module. Setting all *w*_i_=1 recovers plain anchoring (the anchor present, but its reference unused for weighting); the ablation (Section 6.1) shows the reweighting, which the anchor’s reference makes possible, is what improves on it. The procedure is leakage-safe: the target signal is drawn only from its training fold (recomputed per cross-validation fold), and the held-out test fold never enters pre-training, fine-tuning, or model selection. Molecules of the test fold may, however, appear in upstream datasets measured under other conditions, so the setting is condition-held-out with molecules transductive. We use the first-order approximation (stop-gradient through the inner update), which is inexpensive because the adapted set (the head and the optional context) is small.

#### Algorithm 1

Target-Informed Pretraining

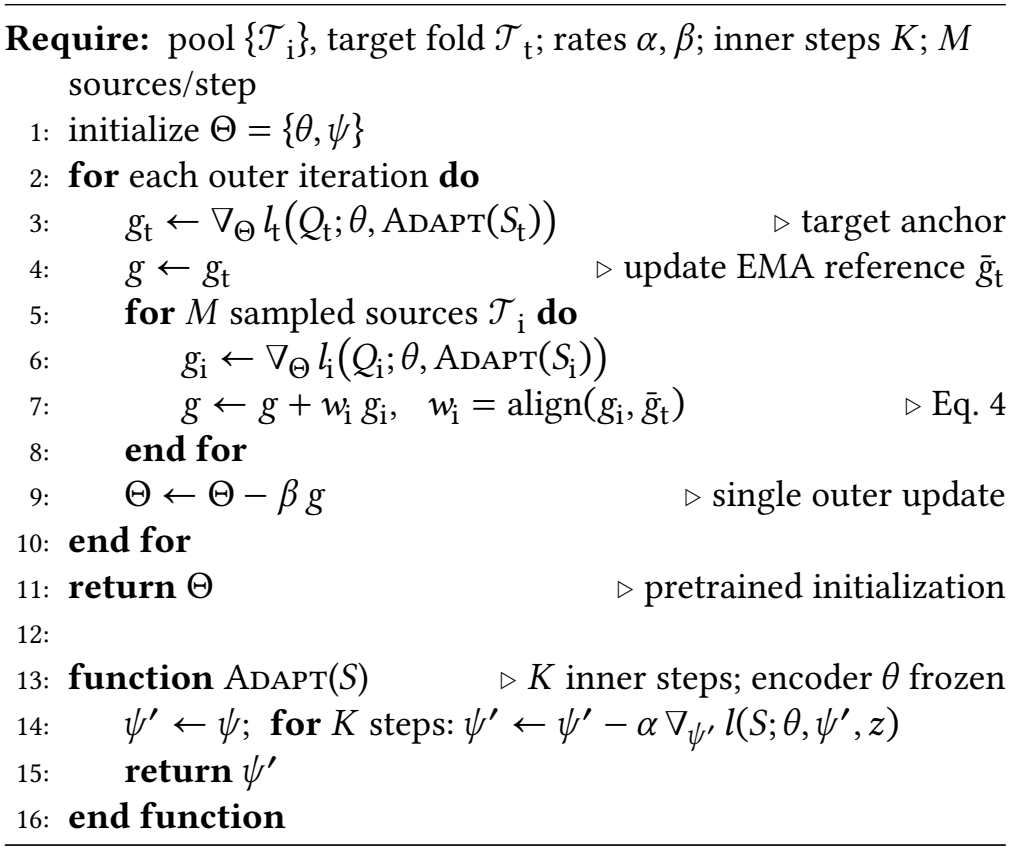

### 3.2 Hyperparameter Selection

We set the inner learning rate *α* = 0.001 and outer rate *β* = 0.0001 empirically, consistent with standard meta-learning practice [11, 24]: a small inner step and a ten-fold smaller outer step (*β* < *α*) keep meta-updates stable while still adapting to the target, taking *K* = 5 inner-loop steps on support sets of *k* = 16 examples. First-order meta-learning theory [10] likewise favors a small inner rate for convergence; since our objective is non-convex and the theory’s assumptions are hard to verify, we treat this as guidance rather than a guarantee. The sweep in Appendix B.1 confirms the configuration is near-optimal.

## 4 Experimental Setup

We evaluate TIMPANI on the RepoRT database under a leakage-safe 5-fold cross-validation protocol. On our primary, metadata-free backbone we compare TIMPANI against TL, and we further test a metadata-conditioned graph transformer and four recent external RT baselines.

### Datasets and upstream pool

We curate the RepoRT database (421 datasets, each with chromatographic-condition metadata) by keeping small molecules (≤ 300 atoms) over eleven elements (C, O, N, H, P, S, F, Cl, B, Br, I), averaging repeated measurements per compound, deduplicating by canonical SMILES (a leakage control), and removing non-eluting compounds below a per-dataset void thresh-old (the co-elution plateau at the shortest retention times). The 38 datasets with more than 100 eluting compounds (34 RP, 4 HILIC) are screened under TL (Section 2). Collapsing two near-duplicate TL-difficult pairs (merging report 0184 into 0183 and 0383 into 0382) leaves 36 (33 RP, 3 HILIC), all of which are evaluated (Section 5.1), with ten TL-difficult targets analyzed in detail.

For each target, TIMPANI’s upstream pool is mode-matched, comprising the same-mode RepoRT datasets large enough to form disjoint support/query sets, excluding the target and its same-condition replicates (datasets that re-measure the target’s compounds under the same condition, identified from the LC metadata). Datasets sharing compounds but acquired under a different condition are kept as legitimate upstream tasks. This yields roughly 190 reversed-phase and 28 HILIC upstream datasets per target, and the target’s own dataset never enters the pool, although 68% of its test-fold compounds (71% counting Tanimoto ≥ 0.7 neighbors) are measured upstream under other conditions (Section 3.1).

### Data splitting and leakage control

All results use 5-fold cross-validation with a fixed split seed (42), training on four folds and testing on the fifth each round, with a random 10% of the training set held out for early stopping during fine-tuning. Upstream datasets are split the same way (training portion for pre-training, held-out portion for validation). Leakage control is central. No compound from a target’s test fold enters fine-tuning or model selection, and the target’s dataset is excluded from pre-training. The evaluation is therefore condition-held-out with molecules transductive. Compound- and scaffold-disjoint results are presented in Section 5.2.

### Backbones

Our primary backbone is the GIN [30], which Kwon et al. [20] found best among GNNs for RT prediction on most Re-poRT datasets. We build molecular graphs from isomeric SMILES with RDKit [28], using its standard atom and bond features. To test whether the gains depend on architecture or metadata, we also evaluate TIMPANI on Graphormer-RT [27], a graph transformer that ingests chromatographic metadata, comparing TL and TIMPANI on the same metadata-aware architecture to isolate the training strategy. TL pre-trains on a single large upstream dataset (METLIN-SMRT for GIN [20]; the mode-matched RepoRT pool for Graphormer-RT, per its published recipe [27]), whereas TIMPANI meta-learns the initialization over the RepoRT mode-pool. Because TL and TIMPANI thus differ in upstream data, we isolate target anchoring through an ablation that holds the pool fixed (Section 6.1).

### Loss, implementation, and reproducibility

We train with the Huber loss [17] (*δ* = 1), sensitive to small time differences yet robust to the outliers common in retention data, and optimize with Adam. Training runs target-informed pre-training followed by frozen-encoder fine-tuning, with TL matched to the same settings. Networks use PyTorch and the Deep Graph Library on a single consumer GPU (11 GB). Pre-training is a one-time offline cost (about one hour per model for TL, half an hour for TIMPANI). All runs fix the primary seed 42 for splits, initialization, and optimization. Robustness checks repeat TL and TIMPANI on two fresh fold partitions (split seeds 43 and 44), and the component ablation (Section 6.1) averages three seeds (42–44). Metrics are reported as mean±sd over the five folds. Complete architectures, features, and hyperparameters are in Appendix A.3, with code at https://github.com/josiehong/TIMPANI.

## 5 Results

### 5.1 TIMPANI Outperforms TL on TL-Difficult Datasets

We first examine whether anchoring pre-training on the known target outperforms conventional transfer learning. Using the GIN backbone [30], we compare TIMPANI with conventional TL in an end-to-end evaluation. This comparison intentionally combines two changes: replacing the single-source pre-training dataset (METLIN-SMRT) with a mode-matched source pool, and introducing TIM-PANI’s target anchoring and gradient-based re-weighting. We dis-entangle the contributions of these two factors in the subsequent baseline comparison and ablation study (Sections 5.3 and 6.1). Performance is evaluated using 5-fold cross-validation on all 36 eligible RepoRT datasets and, in detail, on the ten TL-difficult ones.

To assess the general performance of TIMPANI, we begin with all 36 eligible RepoRT datasets (Table 1). Over all 36, TIMPANI is slightly ahead of TL (*R*^2^ 0.837 vs. 0.826), a difference that is not significant (*p* = 0.98), and no worse than TL beyond a margin of 0.02 *R*^2^ (Wilcoxon non-inferiority *p* < 0.001). Its gains concentrate on the ten TL-difficult datasets (+0.080, 8 of 10 improved, *p* = 0.020) rather than the other 26 (−0.015, *p* = 0.031), and the two groups differ (Mann–Whitney *p* = 0.005), partly reflecting regression to the mean, as the split uses TL’s *R*^2^. Among the difficulty proxies, the gains are largest on HILIC (+0.190), far-condition (+0.052) and small datasets (+0.055), and near zero or negative on their counterparts. All three HILIC targets improve, although with *n* = 3 the paired test cannot reach significance.

On the ten TL-difficult datasets, Table 2 summarizes the results. TIMPANI achieves a higher *R*^2^ than conventional TL on 8 of the 10 target datasets and improves both aggregate metrics, increasing the mean *R*^2^ from 0.60 to 0.68 and reducing the mean MedAE from 0.92 to 0.77 min. These improvements are statistically significant with each dataset as one observation (Wilcoxon signed-rank test on the ten per-target means: *p* = 0.02 for *R*^2^ and *p* = 0.03 for MedAE, sign-flip permutation *p* = 0.016), with a target-clustered bootstrap 95% confidence interval for the mean Δ*R*^2^ of [+0.02, +0.16]. At the individual dataset level, however, a single 5-fold cross-validation provides limited statistical power, as the smallest attainable two-sided Wilcoxon *p*-value is 0.06. To increase power, we repeat TL and TIMPANI on two fresh fold partitions (split seeds 43 and 44), yielding 15 paired folds per target. Under this evaluation, three datasets (RepoRT 0183, 0185, and 0264) improve significantly after Holm correction (marked in Table 2), while 0027 and 0420 reach *p* < 0.05 only without correction. These *p*-values are approximate, as the three partitions reuse the same compounds.

**Table 2:** GIN benchmark on the ten TL-difficult datasets (5-fold CV, seed 42): MedAE (min) and *R*^2^ (mean ± std) for TL vs. TIMPANI, grouped by mode. Bold: best *R*^2^ per row (both metrics in the Average row). ^*^: TIMPANI > TL after Holm correction (per-target Wilcoxon over 15 folds, seeds 42–44).

|  | TL |  | TIMPANI |  |
|---|---|---|---|---|
| ID | MedAE | *R*<sup>2</sup> | MedAE | *R*<sup>2</sup> |
| *HILIC* |  |  |  |  |
| 0027 | 1.35±0.27 | 0.24±0.31 | 1.30±0.22 | **0.50±0.22** |
| 0183 | 1.46±0.47 | 0.33±0.25 | 0.96±0.37 | **0.56±0.23**<sup>\*</sup> |
| 0185 | 0.75±0.15 | 0.75±0.05 | 0.52±0.13 | **0.82±0.07**<sup>\*</sup> |
| *RP* |  |  |  |  |
| 0180 | 0.61±0.15 | 0.72±0.18 | 0.42±0.16 | **0.80±0.17** |
| 0262 | 0.77±0.11 | **0.80±0.12** | 0.79±0.07 | 0.79±0.11 |
| 0263 | 0.91±0.19 | **0.74±0.09** | 0.96±0.23 | 0.70±0.05 |
| 0264 | 0.49±0.15 | 0.69±0.13 | 0.41±0.10 | **0.76±0.05**<sup>\*</sup> |
| 0382 | 0.68±0.07 | 0.76±0.12 | 0.70±0.09 | **0.77±0.08** |
| 0401 | 0.25±0.05 | 0.72±0.08 | 0.20±0.07 | **0.78±0.06** |
| 0420 | 1.92±0.50 | 0.22±0.19 | 1.44±0.37 | **0.28±0.17** |
| Average | 0.92 | 0.60 | **0.77** | **0.68** |

The largest improvements are observed on the HILIC targets, where *R*^2^ increases from 0.24 to 0.50 for RepoRT 0027 and from 0.33 to 0.56 for RepoRT 0183, and TIMPANI raises RepoRT 0185 above the practical threshold of *R*^2^ ≥ 0.8. On the remaining targets, improvements are generally positive but fall within statistical variation, while TIMPANI performs only slightly worse than conventional TL on RepoRT 0262 and 0263. Performance remains modest on RepoRT 0183 and 0420 across all methods, but this does not indicate a limitation of TIMPANI’s transfer strategy. A within-dataset descriptor-based random forest trained without transfer, with full access to the target compounds, achieves only approximately *R*^2^ = 0.50 and 0.24 (Appendix B.2).

### 5.2 Robustness to Fold Partitions, Compound Overlap, and Scaffolds

Table 3 repeats the comparison on the two fresh fold partitions of Section 5.1, with every test-fold molecule removed from TIMPANI’s upstream pool (compound-disjoint pool, TL unchanged), and with folds grouped by Bemis–Murcko scaffold. The gain holds on every fresh partition. Without the test molecules in the pool, TIMPANI loses only 0.024 *R*^2^ relative to the full pool (*p* = 0.19) and stays ahead of TL on each partition, significantly only on seed 44 (mean over seeds, *p* = 0.11). Under the scaffold split, TL collapses on all three partitions (negative *R*^2^ on HILIC targets 0027 and 0183), while TIMPANI keeps most of its accuracy (mean *R*^2^ 0.571 vs. 0.666 on random splits) and stays significantly ahead of TL (*p* = 0.027).

**Table 3:** Robustness on the ten TL-difficult targets (GIN, 5-fold CV, mean *R*^2^). Seed: fold-partition seed. p: paired Wilcoxon over the ten targets (mean rows: each target first averaged over seeds 42–44).

| Setting | TL | TIMPANI | Δ*R*<sup>2</sup> | *p* |
|---|---|---|---|---|
| *Random split* |  |  |  |  |
| Seed 42 (Table 2) | 0.597 | 0.676 | +0.080 | 0.020 |
| Seed 43 | 0.577 | 0.659 | +0.082 | 0.037 |
| Seed 44 | 0.516 | 0.664 | +0.148 | 0.020 |
| **Mean, seeds 42–44** | **0.563** | **0.666** | **+0.103** | **0.014** |
| *Compound-disjoint pool* |  |  |  |  |
| Seed 42 | 0.597 | 0.652 | +0.055 | 0.32 |
| Seed 43 | 0.577 | 0.625 | +0.048 | 0.16 |
| Seed 44 | 0.516 | 0.647 | +0.131 | 0.037 |
| **Mean, seeds 42–44** | **0.563** | **0.641** | **+0.078** | **0.11** |
| *Scaffold-disjoint split* |  |  |  |  |
| Seed 42 | 0.238 | 0.588 | +0.350 | 0.049 |
| Seed 43 | 0.293 | 0.557 | +0.264 | 0.064 |
| Seed 44 | 0.339 | 0.568 | +0.229 | 0.020 |
| **Mean, seeds 42–44** | **0.290** | **0.571** | **+0.281** | **0.027** |

### 5.3 Comparison with the Metadata-Aware Baseline

We next compare TIMPANI with four recent RT prediction methods (DNNpwa [18], MDC-ANN [29], MDL-TL [22], and Graphormer-RT [27]). The first three fail to converge consistently on the low-data cross-validation folds, leaving the metadata-aware Graphormer-RT as the only stable external baseline and therefore our primary point of comparison.

GIN+TIMPANI performs comparably to the metadata-aware Graphormer-RT (mean *R*^2^ 0.68 vs. 0.70, MedAE 0.77 vs. 0.73 min; Wilcoxon signed-rank test *p* = 0.77) while using no per-dataset chromatographic metadata and fine-tuning approximately 34× faster (4 vs. 138 s per fold). A complete comparison with all four baselines, together with TIMPANI applied to the Graphormer-RT backbone, is provided in Appendix B.2. That variant activates TIM-PANI’s optional metadata-conditioning module (Appendix A.3), yet its target-informed components provide no additional benefit beyond the recorded metadata (*R*^2^ 0.68, MedAE 0.72 min, *p* = 0.63).

TIMPANI thus does not improve on Graphormer-RT. It helps most when the backbone has no metadata channel.

To understand this result, we ask how much Graphormer-RT’s accuracy depends on the metadata that a metadata-free model never sees, which real datasets often lack (22% of RepoRT datasets miss a load-bearing condition field and 41% miss temperature; per-field rates in Appendix A.1). At fine-tuning and evaluation, we replace each of the eleven recorded condition fields with the default value the model receives when a field is genuinely absent, and re-measure Graphormer-RT with TL. We contrast the clean model (all fields present) with full removal (every field missing) and with resampling, which replaces each of the target’s condition values by another drawn at random, with replacement, from the same record. Metadata-free GIN+TIMPANI is a fixed reference, and the pool’s metadata is left intact, as for a new target whose own method is undocumented.

Removing all recorded metadata barely affects Graphormer-RT, lowering mean *R*^2^ only from 0.70 to 0.69 (Table 4), a difference that is not statistically significant (paired Wilcoxon *p* = 0.56, *n* = 10), with no target losing more than 0.09 *R*^2^. The degraded model thus remains not significantly different from both its clean counterpart and metadata-free GIN+TIMPANI (0.68). Two further controls point to the same conclusion. Resampling the target’s metadata does not lower accuracy after fine-tuning (0.71 vs. 0.70, *p* = 0.13), and without target fine-tuning the model with correct metadata reaches a median *R*^2^ of only 0.42 (0.17 with metadata removed, 0.47 with it resampled). We report medians because one target falls to *R*^2^ = −3.19 with correct metadata. Since resampled metadata does as well as correct metadata even without fine-tuning (*p* = 0.49), the metadata channel does not carry the condition on its own. Target fine-tuning thus compensates for the missing metadata, which explains why a metadata-free approach can perform comparably to the metadata-aware model, but it does not show that the model recovers the condition.

**Table 4:** Metadata controls on the ten TL-difficult targets (Graphormer-RT + TL, 5-fold CV, seed 42). The *target’s* recorded metadata is kept (present), replaced by missing-field defaults (removed), or resampled. FT and No FT give *R*^2^ with and without target fine-tuning (mean and median, respectively). GIN+TIMPANI, which uses no metadata, is a reference.

| Method | Metadata | FT | No FT |
|---|---|---|---|
| Graphormer-RT + TL | present | 0.70 | 0.42 |
| Graphormer-RT + TL | removed | 0.69 | 0.17 |
| Graphormer-RT + TL | resampled | 0.71 | 0.47 |
| GIN + TIMPANI | none | 0.68 | – |

## 6 Analysis

### 6.1 Ablation Study

Beyond source-pool pre-training, we evaluate a target-agnostic meta-initialization over the pool of upstream tasks with the target excluded. Unlike pooled empirical risk minimization (ERM), each source dataset is treated as a separate task in the meta-objective rather than concatenated into a single training set. TIMPANI extends this baseline by introducing the target as an anchor: the target training fold participates in every meta-update and provides the reference gradient used to re-weight source tasks through gradient alignment. Anchoring therefore serves as the foundation on which gradient alignment operates, not an independent source of improvement. To isolate the contribution of each component, we evaluate all ten TL-difficult targets (GIN backbone), introducing source pooling, target anchoring, and gradient-alignment re-weighting incrementally while holding the training pipeline otherwise fixed (Table 5).

**Table 5:** Component ablation on the ten TL-difficult datasets (GIN, 5-fold CV, mean *R*^2^ over seeds 42–44). Each column adds one component to the previous one: source-pool pre-training (Pool), the target anchor (+Anchor), and gradient-alignment re-weighting (+Align, full TIMPANI). Bold: best per target. The last row gives mean±SD over targets.

| Mode | Target | Pool | +Anchor | +Align |
|---|---|---|---|---|
| HILIC | 0027 | 0.38 | 0.43 | **0.49** |
|  | 0183 | 0.44 | 0.35 | **0.50** |
|  | 0185 | 0.76 | 0.76 | **0.78** |
| RP | 0180 | 0.76 | 0.76 | **0.83** |
|  | 0262 | 0.78 | **0.80** | **0.80** |
|  | 0263 | 0.67 | **0.68** | **0.68** |
|  | 0264 | 0.73 | 0.71 | **0.76** |
|  | 0382 | 0.72 | 0.70 | **0.77** |
|  | 0401 | 0.70 | 0.70 | **0.78** |
|  | 0420 | 0.12 | 0.12 | **0.29** |
| Mean±SD |  | 0.61±0.22 | 0.60±0.22 | **0.67±0.18** |

**Table 6:**
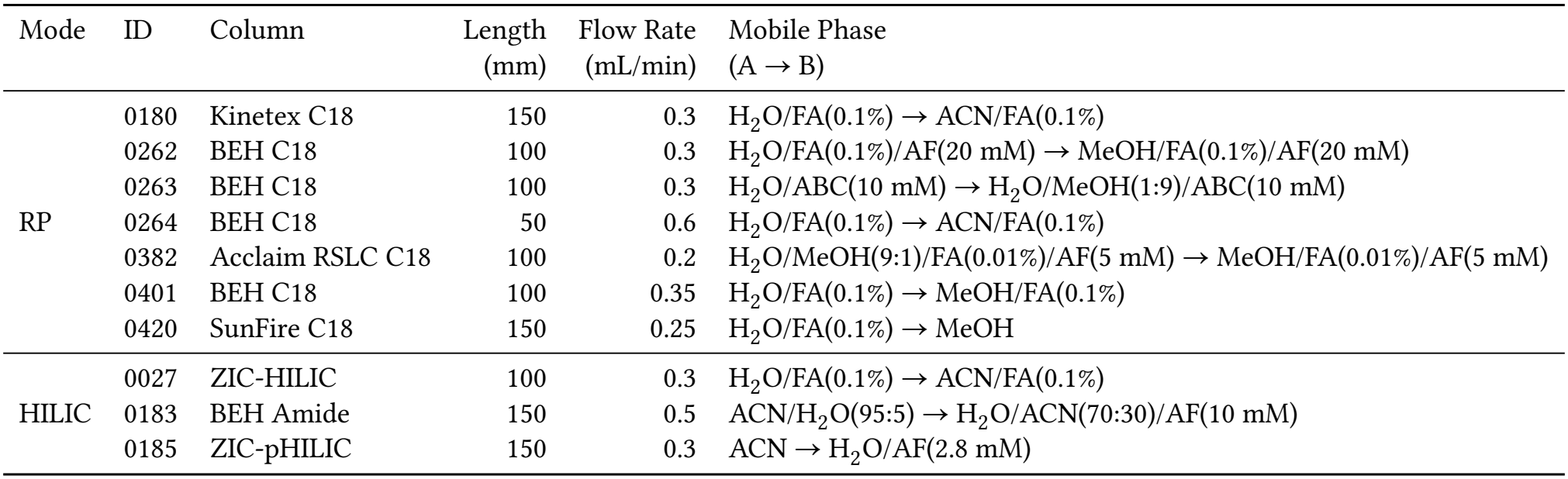
Chromatographic conditions of the ten TL-difficult datasets. FA, AF, and ABC: formic acid, ammonium formate, and ammonium bicarbonate.

| Mode | ID | Column | Length | Flow Rate | Mobile Phase |
|  |  |  | (mm) | (mL/min) | (A → B) |
|---|---|---|---|---|---|
| RP | 0180 | Kinetex C18 | 150 | 0.3 | H<sub>2</sub>O/FA(0.1%) → ACN/FA(0.1%) |
|  | 0262 | BEH C18 | 100 | 0.3 | H<sub>2</sub>O/FA(0.1%)/AF(20 mM) → MeOH/FA(0.1%)/AF(20 mM) |
|  | 0263 | BEH C18 | 100 | 0.3 | H<sub>2</sub>O/ABC(10 mM) → H<sub>2</sub>O/MeOH(1:9)/ABC(10 mM) |
|  | 0264 | BEH C18 | 50 | 0.6 | H<sub>2</sub>O/FA(0.1%) → ACN/FA(0.1%) |
|  | 0382 | Acclaim RSLC C18 | 100 | 0.2 | H<sub>2</sub>O/MeOH(9:1)/FA(0.01%)/AF(5 mM) → MeOH/FA(0.01%)/AF(5 mM) |
|  | 0401 | BEH C18 | 100 | 0.35 | H<sub>2</sub>O/FA(0.1%) → MeOH/FA(0.1%) |
|  | 0420 | SunFire C18 | 150 | 0.25 | H<sub>2</sub>O/FA(0.1%) → MeOH |
| HILIC | 0027 | ZIC-HILIC | 100 | 0.3 | H<sub>2</sub>O/FA(0.1%) → ACN/FA(0.1%) |
|  | 0183 | BEH Amide | 150 | 0.5 | ACN/H<sub>2</sub>O(95:5) → H<sub>2</sub>O/ACN(70:30)/AF(10 mM) |
|  | 0185 | ZIC-pHILIC | 150 | 0.3 | ACN → H<sub>2</sub>O/AF(2.8 mM) |

#### Component contributions

All variants share the source pool and fine-tuning protocol, so differences are attributable to the added component, and each result averages three seeds (42–44). Gradient-alignment re-weighting is the decisive component. Pooling and anchoring alone do not significantly beat single-source TL, and only adding re-weighting yields a consistent, significant gain. Relative to uniform target anchoring, it increases the mean *R*^2^ from 0.60 to 0.67, reduces the mean MedAE from 0.93 to 0.79 min, and improves all ten targets. It is the only variant that significantly beats conventional single-source TL (0.57 to 0.67, *p* = 0.01, with TL averaged over the three seeds rather than its seed-42 value of 0.60 in Table 2), as well as source-pool pre-training and target anchoring (paired Wilcoxon signed-rank test, *p* = 0.002 for both). Source-pool pre-training alone benefits primarily the two most challenging HILIC targets, where conventional TL performs worst, increasing *R*^2^ by 0.24 on RepoRT 0027 and 0.18 on RepoRT 0183, but is otherwise inconsistent, improving only five of the ten targets (mean gain +0.033) and not significantly beating conventional TL (*p* = 1.0). Target anchoring without gradient-based re-weighting likewise provides no measurable benefit over source-pool pre-training (mean *R*^2^ 0.60 vs. 0.61, *p* = 0.85). The target signal thus helps only when used to re-weight sources by gradient alignment. Yet all three components are necessary. The source pool provides the upstream tasks, the target anchor the reference gradient, and gradient-alignment re-weighting combines them into a consistent, statistically significant improvement. Matched controls support this (Appendix B.1): TIMPANI beats pooled ERM, which trains on the same pool plus the target’s training fold without meta-learning, on 9 of 10 targets (0.676 vs. 0.634, *p* = 0.006), TL from the same pool on all ten (0.607, *p* = 0.002), and nearest-source TL with 1 or 5 sources (*p* = 0.037 and 0.002, but *p* = 0.064 with 3). The alignment weights without the meta-learning loop fall below TL (0.568).

#### Meta-training hyperparameters

A one-at-a-time sweep of the remaining meta-training hyperparameters, i.e., the meta-steps and the support-set size *k*, on report 0183 shows that the benchmark configuration is near-optimal and robust across random seeds (Appendix B.1, Table 7).

**Table 7:** Hyperparameter sweep on report 0183 (GIN, 5-fold CV): meta-training steps and support-set size *k*, varied one at a time from the benchmark configuration (^†^). Bold: best *R*^2^ per block.

| Hyperparameter | Value | MedAE | *R*<sup>2</sup> |
|---|---|---|---|
| Meta-training steps | 1000 | 1.29±0.50 | 0.53±0.16 |
|  | 2000 | 1.19±0.52 | 0.51±0.16 |
|  | 4000<sup>†</sup> | 1.01±0.36 | 0.53±0.18 |
|  | 8000 | 0.91±0.12 | **0.59±0.18** |
| Support-set size *k* | 8 | 0.99±0.38 | 0.51±0.19 |
|  | 16<sup>†</sup> | 1.01±0.36 | **0.53±0.18** |
|  | 24 | 1.11±0.46 | 0.48±0.20 |
|  | 32 | 1.11±0.47 | 0.50±0.07 |

### 6.2 The Advantage Emerges as Target Data Grow

We subsampled the target training fold of RepoRT 0183 to increasing sizes (40, 60, 80, and the full ~92-compound fold) and compared SC, TL, and TIMPANI (GIN backbone, 5-fold CV), probing how many labeled target compounds a new condition requires. TIM-PANI consistently converts additional target data into accuracy, increasing *R*^2^ from 0.30 with 40 compounds to 0.52 with the full fold (within experimental variation of the benchmark 0.56) and reducing MedAE from 1.44 to 1.12 min, whereas both TL and SC plateau at approximately *R*^2^ = 0.35 (Figure 3). The advantage emerges once enough target data are available to anchor pre-training, consistent with the ablation.

### 6.3 TIMPANI Starts in a Lower-Loss Basin

To understand why TIMPANI transfers more effectively than conventional TL, we visualized the downstream loss landscape following the procedure of Li et al. [21], perturbing the pretrained parameters along two random filter-normalized directions (Figure 4). Using a shared color scale, the TIMPANI initialization lies substantially lower on the downstream loss surface than the conventional TL initialization, even before fine-tuning: on RepoRT 0183, the center-of-landscape Huber loss is ≈ 0.2 for TIMPANI compared with ≈ 1.7 for TL, a ~8× reduction. These results support the intuition illustrated in Figure 1: target-informed pre-training initializes the model much closer to the target optimum than conventional TL. The same pattern holds on 0185 (HILIC,1.46 to 0.07), 0264 (RP,0.28 to 0.03), and 0401 (RP).

## 7 Related Work

TIMPANI lies at the intersection of three research areas: retention-time prediction under label scarcity, gradient-based meta-learning, and gradient-based task weighting. From these it inherits, respectively, the goal of transferring across chromatographic conditions, the nested inner/outer optimization, and the use of gradient alignment to prioritize source tasks. Its key distinction is a single assumption absent from prior approaches, that the chromatographic condition is fixed and known during pre-training. Whereas existing RT methods pre-train on upstream data alone, MAML-style algorithms target unseen future tasks, and task-weighting methods balance jointly trained tasks, TIMPANI uses the known target to both anchor pre-training and re-weight sources by gradient alignment.

### Retention-time prediction

Beyond the single-condition RT models above, several address cross-condition transfer under limited target labels. DNNpwa [18] pre-trains descriptor networks with weighted autoencoders on SMRT. MDC-ANN [29] selects target-specific combinations of molecular representations. MDL-TL [22] encodes chromatographic parameters and pre-trains across datasets. Graphormer-RT [27] conditions on chromatographic metadata and trains jointly across many RepoRT datasets. Despite these differences, all pre-train on upstream data alone.

### Meta-learning

TIMPANI’s optimization framework is closely related to gradient-based meta-learning methods such as MAML [1, 11], Reptile [24], and ANIL [25], which learn an initialization that adapts rapidly to tasks sampled from a distribution. TIMPANI differs in objective and deployment. Rather than optimizing for an unknown future task, it optimizes toward a single known target, whose training data define the inner update and provide the reference gradient for every outer update. At deployment, the meta-learned encoder stays frozen and only the prediction head is fine-tuned. The objective is therefore not a generally adaptable initialization but one tailored to a target condition seen during training.

### Gradient-based task weighting

TIMPANI’s source re-weighting, which scales each source task by the cosine similarity between its gradient and the target gradient, is related to gradient-based multitask optimization methods such as PCGrad [32], GradNorm [4], gradient-similarity auxiliary-task weighting [9], and transfer-affinity task grouping [26]. These methods estimate relationships among jointly optimized tasks. TIMPANI instead measures each upstream task’s alignment with a single fixed target and uses it to steer meta-learning toward that target. Because the target is available throughout training, the setting is closer to transductive transfer or domain adaptation [12] than to the inductive setting of adapting to unseen tasks.

## 8 Conclusion and Future Work

We presented TIMPANI (Target-Informed Pretraining), a frame-work that meta-learns a target-specific initialization over a pool of chromatographic conditions, using the target’s own training data to anchor pre-training and re-weight source tasks through gradient alignment, followed by fine-tuning with the encoder frozen. Across ten TL-difficult RepoRT datasets, TIMPANI significantly outper-forms conventional TL (mean *R*^2^ 0.68 vs. 0.60, *p* = 0.02), while three of four recent RT prediction methods fail to converge in this low-data regime. Ablation studies identify target-informed gradient re-weighting as the primary source of these gains, while a loss-landscape analysis shows that TIMPANI initializes substantially closer to the downstream optimum than conventional TL. Without per-dataset chromatographic metadata, TIMPANI performs comparably to the metadata-aware Graphormer-RT. Our controls attribute this to target fine-tuning, which compensates for missing metadata. These results demonstrate that accurate condition-specific RT prediction can be achieved from a small in-house dataset augmented with publicly available data.

More broadly, TIMPANI is applicable whenever labels depend jointly on molecular structure and a known but data-scarce experimental condition whose structure–property relationship is condition-specific, as in retention-time prediction but not size-dominated collision cross-section prediction. It is especially attractive when the condition cannot be meaningfully encoded as metadata. In chiral chromatography [15], for instance, enantioselectivity can reverse the elution order across stationary phases, leaving metadata-conditioned models little to condition on, whereas TIM-PANI adapts to it through the target data (given a stereochemistry-aware molecular representation). Extending TIMPANI to such settings, and predicting condition compatibility for source selection without shared compounds, are left for future work.

## Acknowledgments

We are grateful to Drs. Christopher J. Welch, Chris Brown, Rongrong Dai, and David J. Wild for their insightful suggestions. We acknowledge the Center for Bioanalytical Metrology (CBM), an NSF Industry-University Cooperative Research Center, for providing funding under grant NSF IIP-2413195. This work was also partially supported by National Science Foundation grant DBI-2011271.

## Limitations and Ethical Considerations

### Limitations

TIMPANI learns a separate target-specific initialization for each chromatographic condition, incurring a one-time offline pre-training cost per new target, and requires both a small labeled target dataset and a pool of same-mode source datasets. Its benefits are confined to problems where the structure–property relationship depends strongly on the experimental condition, and the gains diminish when chromatographic metadata already capture the condition effectively, or on datasets that are hard for structure-based prediction in general, as observed for RepoRT 0183 and 0420. Our evaluation covers only RT prediction on RepoRT and METLIN-SMRT. We have no external or prospective dataset to test on, and we did not analyze what the learned representations encode. The frame-work should extend to other condition-dependent tasks, though this remains to be demonstrated.

### Ethical considerations

We use only public datasets (RepoRT and METLIN-SMRT) with no human subjects or personal information, so the work raises no privacy concerns and needed no institutional review. Its impact is chiefly beneficial, lowering the data and cost barrier to compound identification for resource-limited labs, with no specific misuse foreseen beyond existing chromatographic work-flows. One caveat is bias, as RepoRT skews toward reversed-phase methods and common chemical classes, so predictions may be less reliable for under-represented conditions and should be validated on the user’s own instrument before deployment.

## Generative AI Usage

Per the ACM and KDD 2027 policies, we used a large language model (Anthropic’s Claude, via Claude Code) to polish the writing and to assist with debugging and boilerplate in the experiment, analysis, and figure code. It did not conceive the method, design the experiments, generate results, or produce citations, and is not an author; the authors wrote the core implementation, verified all content, and take full responsibility.

## A Datasets and Implementation Details

### A.1 Dataset Scale and Experimental Conditions

Most RepoRT conditions are data-scarce: of the 421 datasets, the median holds only 83 compounds and 56% fall below 100, while a single dataset, METLIN-SMRT (80,038 compounds), dwarfs the rest and is the lone large source that TL relies on.

Table 6 lists the key chromatographic conditions of the ten TL-difficult datasets, which vary widely. Closely related datasets can still differ in ways that matter: report 0262 and 0263 share the same column and dimensions but use different buffers (formic acid with ammonium formate vs. ammonium bicarbonate).

RepoRT metadata is also frequently incomplete. Across all 421 datasets, the fraction missing each condition field is: temperature 41.1%, eluent B pH 11.9%, eluent A pH 11.4%, flow rate 10.0%, column length and inner diameter 6.9% each, particle size 6.7%, and column manufacturer and USP code 2.1% each.

### A.2 Condition Distance and Molecular Overlap

#### Chromatographic-condition distance

Each LC condition is featurized by seven numeric fields from the RepoRT metadata and gradient table (column length, particle size, inner diameter, flow rate, and the starting %B, maximum %B, and duration of the gradient), median-imputed and standardized. The distance is the Euclidean distance between these vectors plus +2 for different modes (RP vs. HILIC) and +1 for different USP column codes. Figure 2b and e use it. Temperature is excluded because 41% of datasets lack it, and including it leaves the correlations unchanged.

#### Molecular overlap

The fraction of target compounds with at least one source compound at Tanimoto similarity ≥ 0.7 (Morgan finger-prints, radius 2, 2048 bits). Figure 2c measures it against METLIN-SMRT.

### A.3 Architectures and Hyperparameters

#### Backbones

The GIN has five layers (embedding dimension 300, dropout 0.1), mean pooling, and a two-layer MLP head (hidden 256). Atoms carry 41-dimensional one-hot features (element, formal charge, degree, hybridization, hydrogen count, valence, chirality, ring size, aromaticity), and bonds 8-dimensional ones (type, stere-ochemistry, ring and conjugation flags). Graphormer-RT follows Stienstra et al. [27]: eight layers, embedding dimension 512, 64 heads, attention dropout 0.15, a four-layer MLP head, and meta-data encoded by two 256-hidden encoders summed into a 512-dimensional condition node.

#### Meta-training (TIMPANI)

Inner learning rate *α* = 10^−3^, outer (meta) learning rate *β* = 10^−4^, *K* = 5 inner gradient steps, support and query sets of *k* = 16 compounds each, 4000 meta-steps, and *M* = 4 source tasks accumulated per step alongside the target. Each pool dataset contributes at most 100 compounds per meta-step. Fine-tuning freezes the encoder and trains the head (learning rate 5×10^−5^, up to 150 epochs) with early stopping (patience 20, 10% validation split).

#### Optional condition context

The primary GIN backbone is metadata-free: the context is omitted (*z* = *∅*) and the inner loop adapts the head alone, so the condition module is optional, not a prerequisite. On the metadata-aware Graphormer-RT backbone (Appendix B.2) it is active: when chromatographic metadata *c* are recorded, we initialize the context from them, *z*^0^ = *g*_*ϕ*_(*c*), where *g*_*ϕ*_ is a small encoder with a learned token for missing fields. We add *z* to the inner-loop adapted set, *z*^′^ = *z* − *α∇*_*z*_*l*(*S*; ·). The encoder *g*_*ϕ*_ is meta-learned in the outer loop.

#### Void threshold and pool construction

Non-eluting compounds are removed per dataset below a void threshold, the last co-elution plateau (a repeated RT) before three consecutive unique RTs (0 for well-separated datasets). Each target’s pool holds every same-mode RepoRT dataset with at least *k*_support_ + *k*_query_ = 32 eluting compounds, excluding the target and its same-condition replicates.

## B Additional Results and Analyses

### B.1 Hyperparameter Sensitivity and Matched Controls

In the sweep on report 0183 (Table 7), meta-steps 8000 scores highest but does not survive a multi-seed check (Section 6.1), and no support-set size *k* beats 16. Table 8 compares TIMPANI with controls that hold the source data or the target supervision fixed. Target-only training uses the target’s training fold alone. Pool TL pre-trains on TIMPANI’s pool with the same step budget. Pooled ERM adds the target’s training fold to the pool, without meta-learning or alignment. Nearest-source TL pre-trains on the 1, 3, or 5 datasets closest in LC condition, and alignment without meta-learning applies TIM-PANI’s weights in ordinary training.

**Table 8:** Matched controls on the ten TL-difficult targets (GIN, 5-fold CV, seed 42): mean *R*^2^, TIMPANI wins (targets where TIMPANI has the higher *R*^2^), and paired Wilcoxon *p* of TIM-PANI vs. the control on per-target means. Nearest-source TL: best of 1, 3, or 5 sources (all in Section 6.1).

| Method | *R*<sup>2</sup> | TIMPANI<br>wins | *p* |
|---|---|---|---|
| Target-only training | 0.479 | 10/10 | 0.002 |
| TL from METLIN-SMRT | 0.597 | 8/10 | 0.020 |
| Pool TL (TIMPANI’s pool) | 0.607 | 10/10 | 0.002 |
| Pooled ERM (pool + target fold) | 0.634 | 9/10 | 0.006 |
| Nearest-source TL (3 sources) | 0.603 | 8/10 | 0.064 |
| Alignment without meta-learning | 0.568 | 10/10 | 0.002 |
| TIMPANI | 0.676 | – | – |

### B.2 Full Baseline Comparison

Table 9 compares TIMPANI against the four external RT base-lines [18, 22, 27, 29], each run with its authors’ published configura-tion on identical folds. The three non-metadata baselines (DNNpwa, MDC-ANN, MDL-TL) diverge on many low-data folds (*R*^2^ < 0 on 16, 37, and 12 of 50 folds), so MedAE is the primary metric. TIMPANI on Graphormer-RT (Ours) and Graphormer-RT with TL are not significantly different (*p* = 0.63). The random forest of Section 5.1 uses 300 trees on RDKit 2D descriptors, trained on the same folds without upstream data.

**Table 9:** Comparison with four external RT baselines on the ten TL-difficult datasets (5-fold CV, seed 42): MedAE (min) and *R*^2^, mean ± std. ↓/↑: lower/higher is better. Ours is Graphormer-RT with TIMPANI, and Graphormer-RT is the same backbone with TL. Bold/underline: best/second best per row. ≪ 0: *R*^2^ far below zero, dominated by a divergent fold. Superscript: number of folds (of 5) with *R*^2^ < 0, in total 16, 12, 37, and 2 of 50 for DNNpwa, MDL-TL, MDC-ANN, and Graphormer-RT.

| Metric | Mode | ID | DNNpwa | MDL-TL | MDC-ANN | Graphormer-RT | Ours |
|---|---|---|---|---|---|---|---|
| MedAE↓ | HILIC | 0027 | 1.53±0.57 | 1.80±0.65 | 7.08±6.75 | **0.72±0.20** | 0.75±0.27 |
|  |  | 0183 | 3.37±1.45 | 3.12±0.99 | 7.46±4.89 | 0.86±0.33 | **0.84±0.28** |
|  |  | 0185 | 0.99±0.15 | 1.00±0.22 | 4.07±2.14 | 0.38±0.10 | **0.33±0.07** |
|  | RP | 0180 | 0.51±0.17 | 0.50±0.07 | 1.60±0.93 | **0.14±0.04** | 0.34±0.17 |
|  |  | 0262 | 1.15±0.17 | 1.70±0.24 | 3.25±0.50 | 0.94±0.12 | **0.88±0.09** |
|  |  | 0263 | 1.52±0.33 | 1.52±0.43 | 3.27±2.06 | 1.03±0.10 | **0.91±0.09** |
|  |  | 0264 | 0.72±0.07 | 0.69±0.13 | 1.68±0.74 | 0.46±0.08 | **0.35±0.03** |
|  |  | 0382 | 2.38±0.82 | 1.20±0.08 | 4.01±1.27 | 0.82±0.13 | **0.80±0.08** |
|  |  | 0401 | 0.48±0.07 | 0.39±0.08 | 1.23±0.23 | 0.23±0.08 | **0.18±0.06** |
|  |  | 0420 | 2.15±0.65 | **1.56±0.28** | 2.29±0.49 | 1.68±0.54 | 1.80±0.33 |
| *R*<sup>2</sup> ↑ | HILIC | 0027 | 0.18±0.28<sup>(1)</sup> | −0.17±0.29<sup>(4)</sup> | ≪ 0<sup>(4)</sup> | 0.77±0.09 | **0.81±0.09** |
|  |  | 0183 | ≪ 0<sup>(4)</sup> | ≪ 0<sup>(3)</sup> | ≪ 0<sup>(5)</sup> | **0.47±0.28**<sup>(1)</sup> | 0.47±0.29 |
|  |  | 0185 | ≪ 0<sup>(1)</sup> | ≪ 0<sup>(1)</sup> | ≪ 0<sup>(5)</sup> | **0.87±0.03** | 0.84±0.07 |
|  | RP | 0180 | 0.74±0.12 | 0.39±0.45<sup>(1)</sup> | 0.22±0.34<sup>(1)</sup> | **0.89±0.15** | 0.87±0.15 |
|  |  | 0262 | 0.66±0.13 | 0.55±0.15 | −0.84±0.83<sup>(5)</sup> | **0.82±0.11** | 0.80±0.09 |
|  |  | 0263 | 0.48±0.17 | 0.46±0.26 | ≪ 0<sup>(2)</sup> | **0.74±0.05** | 0.74±0.02 |
|  |  | 0264 | ≪ 0<sup>(1)</sup> | 0.61±0.10 | ≪ 0<sup>(5)</sup> | 0.74±0.03 | **0.77±0.05** |
|  |  | 0382 | ≪ 0<sup>(5)</sup> | ≪ 0<sup>(1)</sup> | ≪ 0<sup>(5)</sup> | **0.75±0.09** | 0.70±0.08 |
|  |  | 0401 | 0.68±0.09 | 0.58±0.07 | −0.11±0.42<sup>(2)</sup> | 0.67±0.13 | **0.76±0.08** |
|  |  | 0420 | −0.17±0.52<sup>(4)</sup> | −0.03±0.18<sup>(2)</sup> | −0.09±0.16<sup>(3)</sup> | **0.25±0.30**<sup>(1)</sup> | 0.09±0.38 |

### B.3 Per-Dataset Results on All Eligible Datasets

Figure 5 plots each dataset of Table 1. The gain does not track the percentage of the target’s compounds measured upstream (Spearman *ρ* = −0.11, *p* = 0.56, 31 datasets). On the 21 datasets never used to set the threshold, develop, or tune the method, TIMPANI is equivalent to TL within ±0.02 *R*^2^ (0.914 vs. 0.920, Wilcoxon equivalence test *p* = 0.001). Counting each same-condition replicate cluster once gives +0.029 (*p* = 0.49).

